# Leveraging mouse chromatin data for heritability enrichment informs common disease architecture and reveals cortical layer contributions to schizophrenia

**DOI:** 10.1101/427484

**Authors:** Paul W. Hook, Andrew S. McCallion

**Affiliations:** Department of Genetic Medicine, Johns Hopkins University School of Medicine, Baltimore, Maryland, USA; Department of Comparative and Molecular Pathobiology, Johns Hopkins University School of Medicine, Baltimore, Maryland, USA; Department of Medicine, Johns Hopkins University School of Medicine, Baltimore, Maryland, USA

## Abstract

Genome-wide association studies have implicated thousands of non-coding variants across human phenotypes. However, they cannot directly inform the cellular context in which disease-associated variants act. Here, we use open chromatin profiles from discrete mouse cell populations to address this challenge. We applied stratified linkage disequilibrium score regression and evaluated heritability enrichment in 64 genome-wide association studies, emphasizing schizophrenia. We provide evidence that mouse-derived human open chromatin profiles can serve as powerful proxies for difficult to obtain human cell populations, facilitating the illumination of common disease heritability enrichment across an array of human phenotypes. We demonstrate signatures from discrete subpopulations of cortical excitatory and inhibitory neurons are significantly enriched for schizophrenia heritability with maximal enrichment in discrete cortical layer V excitatory neurons. We also show differences between schizophrenia and bipolar disorder are concentrated in excitatory neurons in layers II-III, IV, V as well as the dentate gyrus. Finally, we use these data to fine-map variants in 177 schizophrenia loci, nominating variants in 104/177 loci, and place them in the cellular context where they may modulate risk.

---

Although genome-wide association studies (GWAS) have implicated thousands of variants in an array of human phenotypes, the variation underlying these signals and cellular contexts in which variants act have remained largely unclear (Visscher et al. 2017). Discernment of disease-relevant variants and cell populations is essential for comprehensive functional investigation of the mechanisms of disease.

Schizophrenia has been robustly investigated through GWAS with the number of associated loci increasing from twelve to 179 independent associations in the last decade (O’Donovan et al. 2008; Pardiñas et al. 2018). However, this increase has not been accompanied by the elucidation of disease mechanisms or an increase in the identification of causal variants. To date, support for mechanisms and/or causal variants have been established for two loci (Sekar et al. 2016; Song et al. 2018). The inability to construct and test mechanisms for schizophrenia largely stems from the inability to separate disease-relevant variants from those in linkage disequilibrium (LD) and from the lack of knowledge about what cells are important for disease risk.

Recent studies have begun to identify cell populations for schizophrenia by leveraging GWAS summary statistics and stratified linkage disequilibrium score regression (S-LDSC) (Finucane et al. 2018; Skene et al. 2018). These studies have focused on human and rodent transcriptional data, with the finest resolution of cell populations provided by mouse single-cell RNA-seq data (scRNA-seq). This approach relies upon the imposition of windows around the transcription start sites of genes with cell-dependent expression. Crucially, mouse data can be leveraged by using corresponding human orthologs. The results from these studies have supported a role for cortical excitatory and inhibitory neurons in schizophrenia risk (Finucane et al. 2018; Skene et al. 2018). However, these studies only capture signal driven by variants residing in selected windows, excluding much of the regulatory landscape. As most variants identified through GWAS occur in non-coding DNA (Maurano et al. 2012), these studies have systematically overlooked the capacity to use these biological signatures in an agnostic manner to construct hypotheses indicting putative, distal cis-regulatory elements.

Ideally, human chromatin data with the same cell population resolution as transcriptome data would be used to provide a regulatory context for variants. However, human chromatin data analyzed with S-LDSC have been limited to easy-to-access cell populations (Ulirsch et al. 2019) or heterogeneous adult tissues, broad cell types, and *in vitro* cell lines (like those data available through the ENCODE consortium) (Finucane et al. 2018; Tansey and Hill 2018; ENCODE Project Consortium 2012; Fullard et al. 2018). Mouse data has the potential to overcome these barriers by providing high resolution of the same populations as scRNA-seq. Recently, mouse single-cell ATAC-seq was used to annotate variants and explore heritability of a variety traits, including schizophrenia (Cusanovich et al. 2018). This study implicated many of the same populations in schizophrenia as previous studies that leveraged expression data. However, which variants are relevant to disease and in which cells those variants may act was not explored. Furthermore, only a limited number of GWAS SNPs are included in S-LDSC analysis and only differentially accessible peaks were analyzed (Cusanovich et al. 2018), limiting the SNPs and regulatory elements for which hypotheses could be generated.

Previously, we have successfully used mouse chromatin data to prioritize common human variants for pigmentation and Parkinson disease (Praetorius et al. 2013; McClymont et al. 2018). In this study, we set out to address whether mouse-derived human open chromatin profiles could be used to prioritize cell populations and variants important to schizophrenia. In this way, data from narrowly-defined cell populations that are inaccessible in humans could be used to provide context for variants and allow for the construction of hypotheses. We evaluate a limited number of strategies for converting mouse open chromatin peaks to human peaks and use heritability enrichment (S-LDSC) to prioritize 27 selected (25 mouse and two human) cell populations across 64 GWAS with an emphasis on schizophrenia. Finally, we combine statistical fine-mapping of variants with mouse-derived human open chromatin data to prioritize variants in schizophrenia loci and predict a cellular context in which those variants may act.

## Results

### A uniform pipeline for processing of mouse ATAC-seq data

We obtained publicly available ATAC-seq data derived from lineage identified cell-types sorted *ex vivo* from mice and from mouse brain single nuclei analyses (Table S1) (Preissl et al. 2018; Matcovitch-Natan et al. 2016; Gray et al. 2017; Mo et al. 2015; Hughes et al. 2017; Hosoya et al. 2018; McClymont et al. 2018). In total, we obtained 25 mouse ATAC-seq datasets encompassing subclasses of six broader cell types (dopaminergic neurons, excitatory neurons, glia, inhibitory neurons, retina cells, and T-cells) (Table S1). These datasets were selected to maximize the range of cell types for analysis, while ensuring inclusion of classes with predicted roles in schizophrenia (Howes and Kapur 2009; Coyle 2006; Nakazawa et al. 2012) and facilitating comparison with single-cell RNA-seq populations analyzed in previous heritability studies (Skene et al. 2018; Finucane et al. 2018).

To compile comparable open chromatin profiles, all ATAC-seq data were processed in an uniform manner. Sequencing for each cell population was aligned to the mouse genome (mm10), replicates were combined, and peak summits were called (see Methods for more details). This resulted in 165,143 summits called per cell type (range: 54,880 to 353,125; median: 130,464) with profiles derived from the single-nuclei data having less summits in general (Table S2).

Recognizing that the variable sequencing depths for the datasets may lead to biases in the number of summits, we sought to obtain peaks with similar levels of evidence. To achieve this, we employed a method used by The Cancer Genome Atlas (Corces et al. 2018) (See Methods). We then added 250 base pairs (bp) to either side of each summit and the uniform peaks were merged within each population. This resulted in 78,115 filtered summits per cell population (range: 38,685 to 119,870) and an average of 62,309 peaks (range: 30,791 to 99,119 peaks)(Table S2).

To ensure that the open chromatin profiles reflected expected cell population identities, read counts for each cell population for the union set of peaks (433,555 peaks) were compared using principal component analysis (PCA) and hierarchical clustering. PCA revealed that the vast majority of variation (70.29%) in the data could be explained by whether the ATACseq data was single-nuclei or bulk not experiment or cell population (Fig. S1A, S1B, S1C). Stepwise quantile normalization and batch correction abolished the variation caused by this technical effect (Fig. S1A).

In general, broad cell types clustered together within hierarchical clustering of correlation (Figure 1A) and when PCA results were projected into two dimensional space (Figure 1B). Only single-cell data from inhibitory medium spiny neurons (“Inhibitory MSN*”) and broad inhibitory neurons (“Inhibitory*”) were separated from the bulk inhibitory neurons in both hierarchical clustering of correlation and t-SNE space (Figure 1A, 1B). Inhibitory MSN neurons clustering with excitatory neurons is consistent with the original analysis (Preissl et al. 2018). This separation could be due to the different tissues and methods used to isolate these cells (from cortex vs. from whole forebrain or sorting vs. single-nuclei) (Preissl et al. 2018; Mo et al. 2015; Gray et al. 2017). Overall, these results establish that the uniformly processed open chromatin profiles appropriately reflect cell-dependent biology.

**Figure 1.**
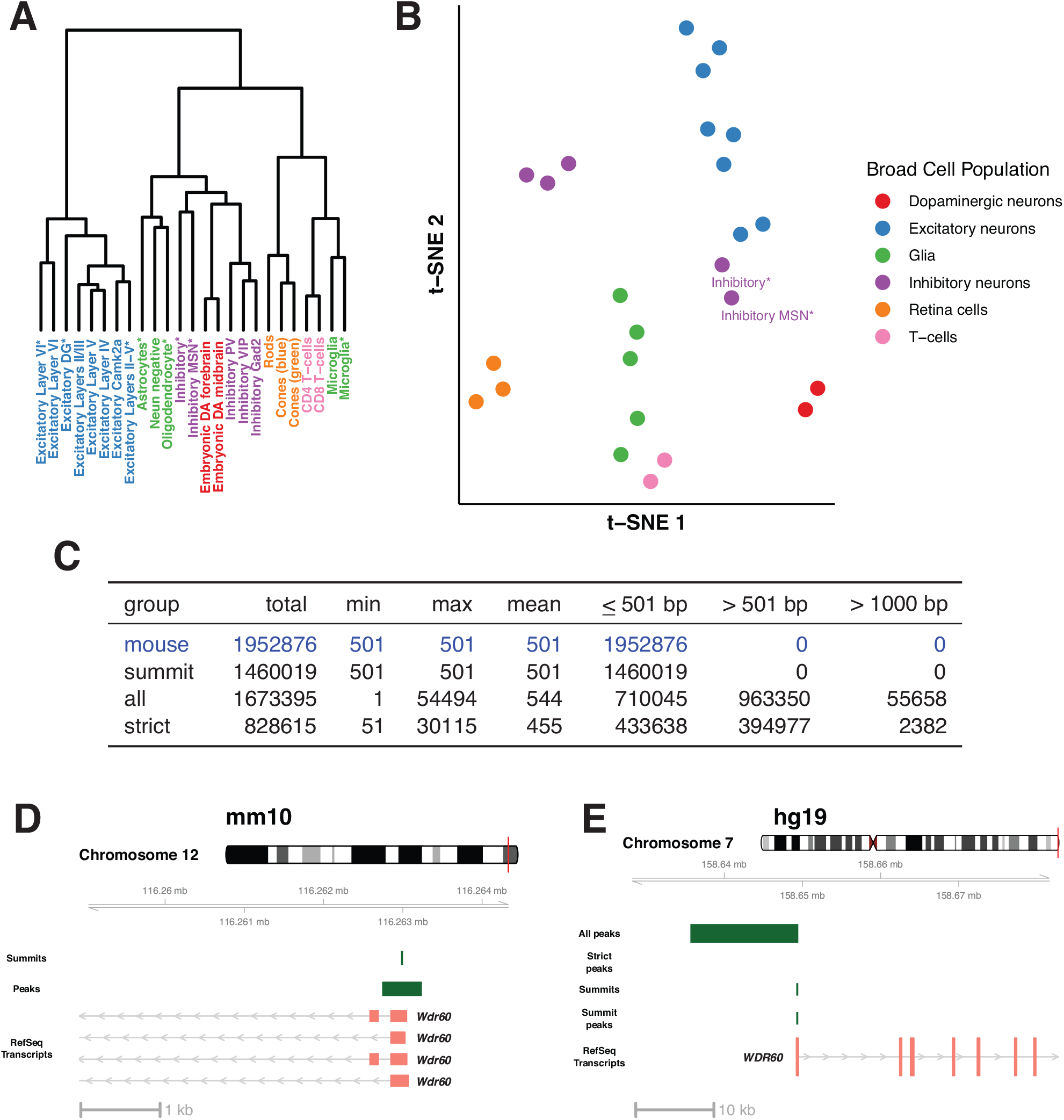
Mouse open chromatin profiles show expected relationships (A, B) and lift over of mouse peaks to human is best done with summits (C, D, E). A) Dendrogram displaying results of hierarchical clustering of the peak count correlations of public, mouse ATAC-seq data. Asterisks in the cell population name indicate single-nuclei datasets. B) t-SNE plot displaying relationships between the peak counts of mouse cell populations. C) Table containing the summary of three lift over strategies applied to public mouse ATAC-seq data. D) Mouse ATAC-seq data at the *Wdr60* promoter region in the mm10 genome. As an example of the data at this locus, summits and peaks from Excitatory Layers II-III are displayed along with RefSeq transcripts. E) Mouse-derived human open chromatin data at the *WDR60* promoter region in the hg19 genome. As an example of data at this locus, lifted over data from Excitatory Layers II-III are displayed along with human RefSeq transcripts. Data includes results from all three lift over strategies employed (“All peaks”, “Strict peaks”, and “Summits”) along with the peak created after summit lift over.

### Converting open chromatin summits from mouse to human provides the most accurate open chromatin profile

We lifted over all open chromatin profiles for these cell populations from the mouse genome (mm10) to syntenic sequences in the human genome (hg19). In order to optimize lift over, we compared three methods. We sought the method which retained the maximum number of peaks while ensuring human profiles resembled mouse profiles. First, peaks were lifted over “as is” including 250 bp extensions with default lift over parameters (“all”). Second, peaks were lifted over “as is” but with much stricter parameters, limiting gap sizes to 20 bp to match previous studies (“strict”)(Vierstra et al. 2014). Finally, we converted the single base pair summits with default parameters and added 250 bp on each side (“summit”).

Although the first method (“all”) resulted in retention of the most peaks from mouse to human (~86%), it also led to a range of peak sizes (1 bp to 54,494 bp) that were vastly different than the uniform input size of 501 bp (Figure 1C). Further, ~58% of lifted over peaks were >501 bp, with ~55,658 peaks doubling in size (>1,000 bp) (Figure 1C). Our second strategy (“strict”) led to ~42% of peaks being lifted over (Figure 1C). This strict parameter did not sufficiently control for peak size with 2382 peaks still exhibiting a peak size greater than 1,000 bp. Finally, the third strategy (“summits”) led to ~75% of peaks being converted while controlling for size (Figure 1C). This third strategy allowed for the mouse-derived human peaks to properly represent mouse peaks. This can be illustrated by observations at the *WDR60* promoter (Figure 1D, 1E). In mouse, one open chromatin summit (from the excitatory neurons from cortical layers II-III) is identified which leads to a peak directly over the *Wdr60* promoter (Figure 1D). When lifted over as a peak, it expands from 501 bp to ~13 kb (“All peaks”, Figure 1E). When strictly controlling for gaps, the peak fails to liftover (“Strict peaks”, Figure 1E). Neither of these results are representative of the regulatory landscape seen in mouse. However, the summit lifts over and produces a 501 bp peak that encompasses the *WDR60* promoter, providing an accurate representation of the data in mouse (“Summits” and “Summit peaks”, Figure 1E). Ultimately, lifting over summits proved the most robust method, retaining a high proportion of peaks while controlling for size and profiles derived from summits were used going forward.

### The majority of mouse-derived human peaks show regulatory activity in human tissues

We next explored how well data from these cell populations recapitulate profiles from orthologous cell populations in humans. Most cell populations included in our study do not have orthologous data generated from humans by design. In order to evaluate the profiles, we compared mouse T-cell profiles (CD4 and CD8) to human open chromatin data from orthologous cell populations processed identically to the mouse data. Additionally, we compared our data to open chromatin data from the Roadmap Epigenome Project (Ernst and Kellis 2015).

We observe 43.5% (16,674/38,299; Figure 2A) of mouse-derived CD8 ATAC-seq peaks overlap with human CD8 ATAC-seq peaks (40,916 peaks) which is slightly higher than previous studies (Vierstra et al. 2014). Using T-cell Roadmap data, we observe 60% (22,927/38,299) and 59% (22,689/38,299) overlap with naive (77,770 peaks) and memory CD8 T-cell data (80,049 peaks) with a slight improvement to 61% (23,698/38,299) when combined (Figure 2A)(90,267 peaks). Further, we find ~83% overlap (31,757/38,299) with peaks found in any Roadmap tissue (493,894 peaks) or the combination of Roadmap and ATAC-seq data (31,796/38,299) (Figure 2A) (624,749 peaks). We observe similar numbers for mouse-derived CD4 T-cells (Table S3).

**Figure 2.**
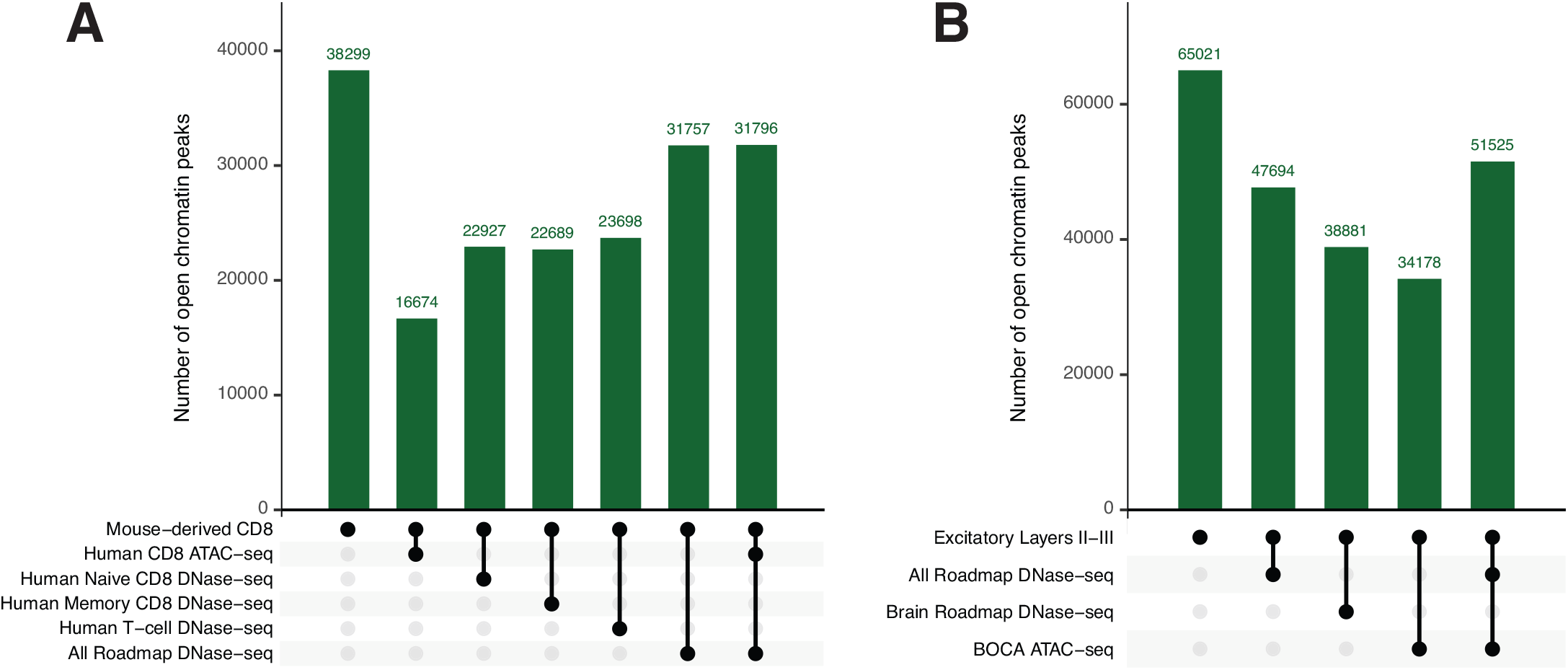
The majority of mouse-derived human open chromatin profiles show regulatory potential in humans. A) Plot displaying the intersection of mouse-derived CD8 T-cell open chromatin peaks with publicly available human datasets. All numbers displayed are the number of mouse-derived peaks that meet the intersecting criteria below the plot. B) Plot displaying the intersection of mouse-derived Excitatory Layers II-III open chromatin peaks with publicly available human datasets. All numbers displayed are the number of mouse-derived peaks that meet the intersecting criteria below the plot.

We extended this analysis to cell populations for which no orthologous data exists. In addition to DNase-seq from Roadmap tissues, we compared profiles to brain-related Roadmap samples only (208,021 peaks) and ATAC-seq data from neurons from the Brain Open Chromatin Atlas (BOCA) (255,977 peaks) (Fullard et al. 2018). As in the T-cell comparisons, all cell populations have the highest overlap with the combination of DNAase-seq and ATAC-seq (Table S4). For example, ~79% of peaks in excitatory neurons in layers II-III (“Excitatory Layers II-III”) overlap with the combined data while we observed 73% with all Roadmap data, 60% with brain-related Roadmap data, and 53% with BOCA data (Figure 2B; Table S4). In summary, the vast majority (average: 78%, range: 70-88%; Table S4) of mouse-derived human open chromatin regions across all cell populations overlap with human open chromatin. This indicates that the large majority of mouse-derived human peaks have regulatory potential in humans and that mouse-derived human peaks are suitable proxies for human cell populations.

### Mouse-derived human profiles recapitulate cell population disease enrichments and reveal new biology

We sought to determine whether mouse-derived chromatin data could be used to inform heritability enrichment of traits and pinpoint cell populations contributing to common phenotypes. We employed S-LDSC to test for enrichment of heritability in open chromatin from 27 cell populations across 64 GWAS. We included open chromatin data from human T-cells to allow for direct comparison to mouse-derived data. The spectrum of traits evaluated included a selection of common neuropsychiatric, neurological, immunological, and behavioral traits, as well as traits from GWAS performed on UK Biobank data (Table S5) (Brainstorm Consortium et al. 2018; Bycroft et al. 2018).

Overall, S-LDSC results for all 64 GWAS established that mouse-derived ATAC-seq data, when lifted over to the human genome, displayed increased heritability enrichment in cell populations consistent with the known biology. In order to explore trait enrichment patterns, we calculated Z-scores of P-values within traits for all S-LDSC results (Table S6). High Z-scores indicate that a cell population has increased heritability for a trait when compared to the other populations. Z-scores were then grouped using hierarchical clustering by cell population and trait. This revealed three broad clusters of cells (Figure 3A, column groups I-III) and three clusters of phenotypes (Figure 3A, row groups A-C). Collectively, these data highlight biological relationships between cell populations and traits (Figure 3A).

**Figure 3.**
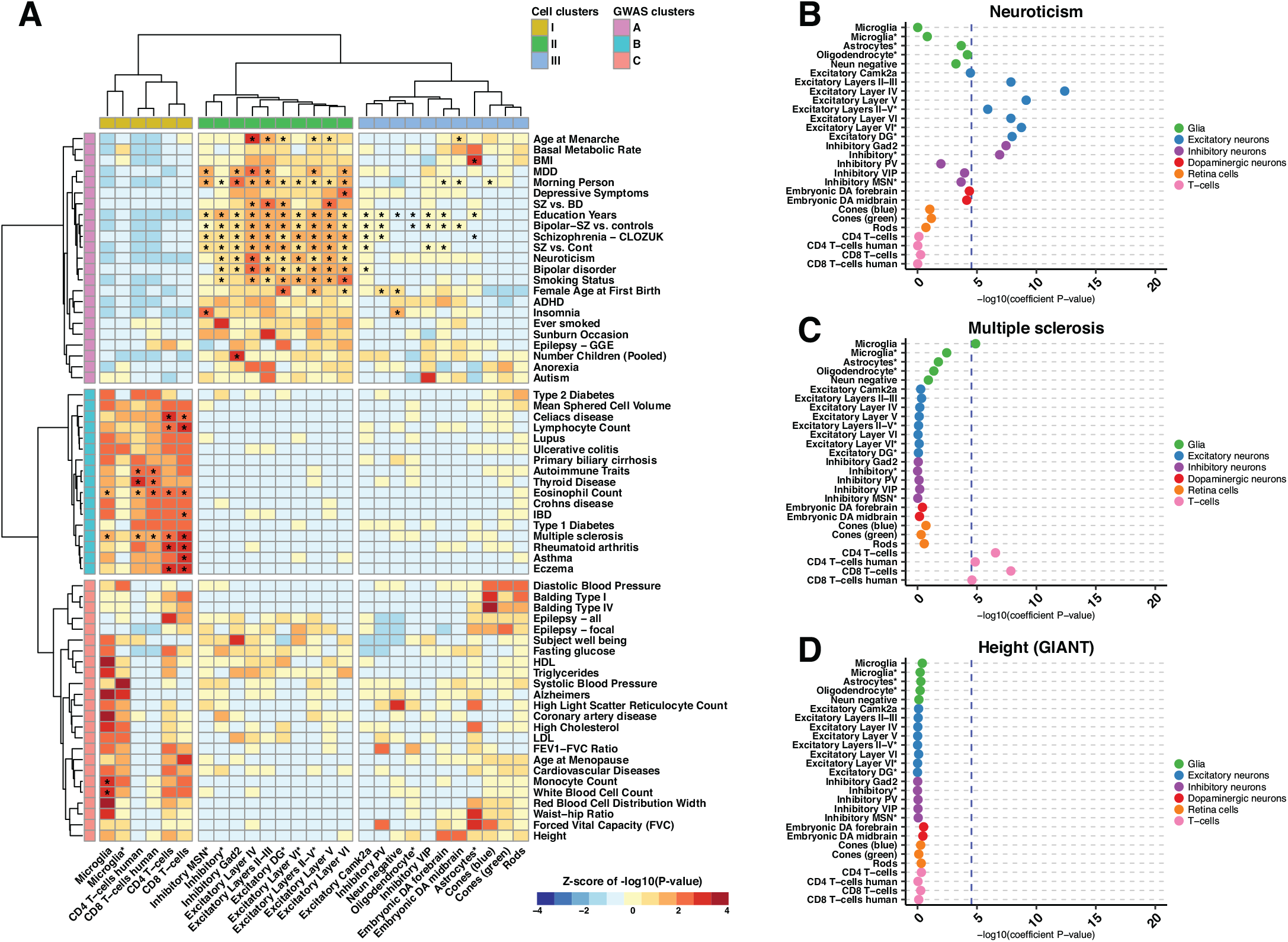
S-LDSC results from 64 GWAS show heritability enrichment in expected cell populations and reveal further insight into disease. A) A heatmap displaying the Z-scores of −log10(P-values) for 27 cell populations across 64 GWAS analyzed. Data was hierarchical clustered by GWAS and cell population. Cell populations that met the across trait significance level (−log10(P) = 4.53857) are indicated with an asterisk. B, C, D) Example dotplots displaying −log10(heritability coefficient P-values) S-LDSC results for GWAS indicative of the observed clustering patterns. Neuroticism (B) for row cluster A, multiple sclerosis (C) for row cluster B, and height from the GIANT consortium (D) for row cluster C. Populations are colored and ordered by broader cell-type category. Asterisks in the cell population name indicate single-nuclei ATAC-seq data. All results can be found in Table S6.

In the first of these highlighted relationships, we observe a collection of broadly defined inhibitory neurons, inhibitory medium spiny neurons (MSNs), excitatory neurons in all cortical layers, and excitatory dentate gyrus (DG) neurons (Figure 3A; row group A, column group II). Cell populations clustering in this group show consistently higher heritability enrichment for neuropsychiatric, neurological, and behavioral phenotypes with many showing significant heritability enrichment (indicated by asterisks) for these traits including neuroticism (Figure 3B). Although perhaps initially surprising, age of menarche, female age at first birth, and number of children are highlighted in this group, cognitive phenotypes display significant genetic correlation with female age at first birth and number of children (Lam et al. 2017).

Grouped together among the adjacent collection of cells are a broadly defined collection of retinal (rods, cones) and nervous system populations (excitatory neurons, glial cells, inhibitory PV and VIP neurons, embryonic dopaminergic neurons) (Figure 3A; row group A, column group 3). While not clustering with the second group of cells, the central nervous system-derived cell populations in this group show enrichment in neurological phenotypes (education years, bipolar disorder, and schizophrenia) that also show enrichment in the second group of cells (Figure 3A, row group A, column group II). However, the enrichments are less than those in the second group. Nonetheless, this group highlights enrichments that may warrant further exploration. The observation that data corresponding to the “morning person” trait reveals enrichment within the blue cone open chromatin region (OCR) profile (Figure 3A) is consistent with the finding that genes expressed highly in retinal tissue are enriched in “morning person” loci (Jones et al. 2019), adding evidence to a potential biological relationship. Additionally, the observed enrichment for astrocyte OCR profile with body mass index heritability (Figure 3A) mirrors mounting evidence that astrocytes and other glia play a role in controlling body weight (García-Cáceres et al. 2012).

The third major grouping (Figure 3A; row group B, column group I) demonstrates heritability for immune-related traits, including lupus, eczema, Crohn’s disease, and general autoimmune traits from the UK Biobank are enriched in open chromatin data from immune cells (microglia, T-cells). Indeed, we detect significant associations for many traits including multiple sclerosis (Figure 3C). Many of these enrichments are consistent with prior data generated using human tissue (Finucane et al. 2018).

By contrast, in the lower portion of this hierarchical clustering analysis (Figure 3A, row group C), the corresponding phenotypes show no such clear biological theme and highlight few cell populations that have enrichments that achieve significance across traits (Figure 3A). It may be expected that traits like height (Figure 3D), fasting glucose, and balding type I, would not reveal significant enrichments in the cell populations we evaluate.

While immune cells show similarly increased heritability enrichment in immune traits, the overlap between traits that reach significance for human T-cells and mouse-derived data is incomplete (Figure 3A). For CD4 T-cells, ≥90% (58/64) of traits are concordant in their enrichment for heritability at a trait-wide level in both human and mouse data. We demonstrate for human CD4 T-cells, four traits were found to reach trait-wide significance (Figure 3A; Table S6). The mouse-derived human CD4 T-cell data shows significance at only 2/4 at the trait-wide level. Additionally, mouse-derived CD4 data reveals four additional traits which reach significance that fail to reach significance using human CD4 data, suggesting that the mouse data may have the power to detect enrichment not yet observed in publicly available human datasets. These general patterns are seen with the CD8 data as well. However, a significance threshold for enrichment is an arbitrary level at which to compare cell populations. In order to explore how mouse-derived human data recapitulated what would be seen in orthologous human data, we compared the heritability regression coefficients between human and mouse T-cells. We observed that in the case of both CD4 T-cells (Spearman’s rho = 0.6005) and CD8 T-cells (Spearman’s rho = 0.6681), the human and mouse-derived data show strong correlation (Figure S2). The observed differences may, in part, reflect the different locations from which the cells were collected (mouse T-cells, thymus; human T-cells, bone marrow/peripheral blood)(Table S1). They may also reflect the power resulting from more homogenous and less challenged immune cell populations that may be obtained from laboratory mice.

Collectively, these results highlight that mouse-derived human profiles broadly recapitulate known biology across a wealth of human phenotypes and thus can serve as suitable proxies for orthologous human cell populations. We also observed that they can illuminate potentially important new biology for a host of traits.

### Schizophrenia heritability is most enriched in cortical layer excitatory neurons

Having established the power of mouse-derived human profiles in studying the genetic architecture of common disease, we restricted our focus to schizophrenia. To facilitate direct comparison with prior transcription-based analyses (Skene et al. 2018), we made use of the recent CLOZUK schizophrenia GWAS (Pardiñas et al. 2018). Of 27 chromatin pro-files, 13 achieved significance when corrected for all traits tested (Figure 4A; Table S6). Our analyses largely indict cortical neurons; with open chromatin profiles from both excitatory and inhibitory populations displaying significant enrichment (Figure 4A; Table S6).

**Figure 4.**
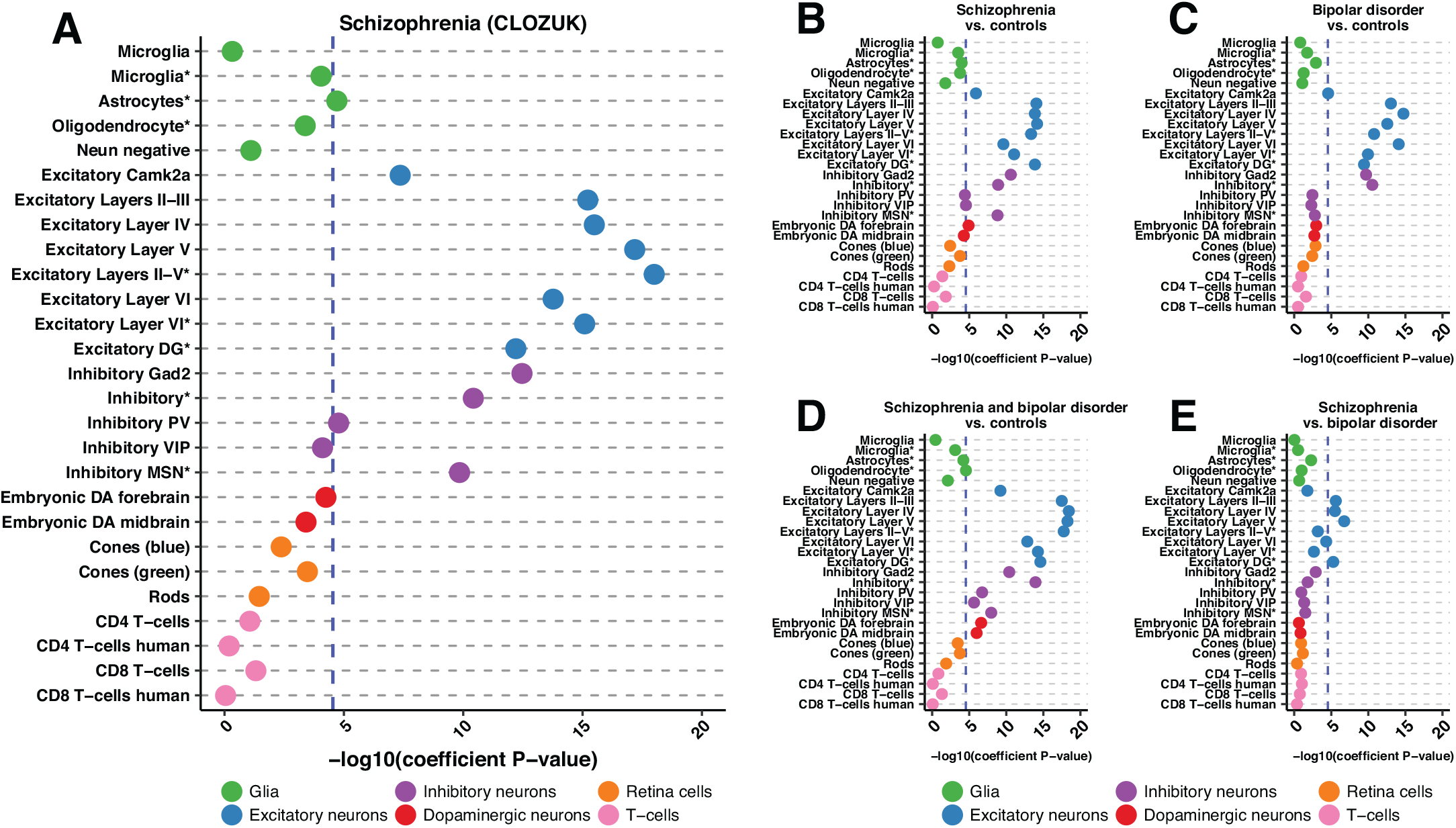
S-LDSC results for CLOZUK and PGC schizophrenia studies as well as bipolar disorder GWAS reveal excitatory cortical neuron enrichment. A, B, C, D, E) Dotplots displaying the −log10(heritability coefficient P-values) S-LDSC results for: A) CLOZUK schizophrenia GWAS, B) PGC schizophrenia GWAS, C) PGC bipolar disorder GWAS, D) schizophrenia and bipolar disorder GWAS and E) PGC schizophrenia versus bipolar disorder GWAS. Across trait significance levels for are shown (-log10(P) = 4.53857; blue dashed line). Populations are colored and ordered by broader cell-type category. Asteriks in the cell population name indicate single-nuclei ATAC-seq data. All results can be found in Table S6.

Within subsets of cortical excitatory neurons, the availability of data from layer-identified populations allowed for detection of a clear and progressive increase in the extent of enrichment, progressing from layers II-III and IV, reaching an apex with open chromatin profiles derived from layer V, and then diminishing slightly in layer VI (Figure 4A; Table S6). This pattern is mirrored in single-nuclei data wherein enrichment in layer III/IV/V cortical excitatory neurons (“Excitatory Layers II-V*”) exceeds that for layer VI cortical excitatory neurons (“Excitatory Layer VI*”) with both being significant (Figure 4; Table S6). Significant enrichment for profiles derived from excitatory neurons of the dentate gyrus (“Excitatory DG*”) provides evidence of additional contribution made by hippocampal excitatory neurons.

Schizophrenia heritability enrichment is not restricted to excitatory neurons; subpopulations of inhibitory neurons also reveal enrichment, with highest levels seen in the broadly defined *Gad2*+ GABAergic population (Figure 4A; Table S6). Notably, enrichment in parvalbumin positive neurons (“Inhibitory PV”) also reaches significance. This inhibitory PV neuron observation echoes a recent study demonstrating that treatment of PV inhibitory neurons in a mouse model of schizophrenia resulted in amelioration of disease phenotypes (Marissal et al. 2018). Beyond the cortex, we detect strong enrichment in *Drd1* positive medium spiny neurons (“Inhibitory MSN*”). Lastly, we note a predicted contribution of glial cells to schizophrenia; chromatin signatures from astrocytes also pass the threshold for significance (Figure 4A; Table S6).

These chromatin based observations are consistent with results using transcriptional data (Skene et al. 2018). These prior data primarily implicated medium spiny neurons, all layers of cortical excitatory neurons, cortical inhibitory neurons, as well as hippocampal CA1 excitatory neurons. We find enrichment in all but hippocampal CA1 excitatory neurons, for which we do not have data. However, we observe enrichment in excitatory neurons derived from the dentate gyrus, which mirror significant schizophrenia enrichment seen in mouse dentate granule cells (Skene et al. 2018). Finally, while astrocytes are not consistently implicated in the transcriptional data, astrocytes from mouse striatum, mouse visual cortex, and human cortex show enrichment for schizophrenia heritability (Skene et al. 2018). Overall, our analysis of open chromatin provides strong orthogonal evidence to transcriptional data for the enrichment of schizophrenia heritability in narrowly-defined cell populations.

### Excitatory neurons in the cortex and hippocampus are enriched for differences between schizophrenia and bipolar disorder

Leveraging our success analyzing schizophrenia, we set out to determine which cell populations may differentiate schizophrenia and bipolar disorder. Although bipolar disorder is related to schizophrenia and their heritabilities are highly correlated, they are unique disorders (Brainstorm Consortium et al. 2018). We took advantage of a recent study that not only performed traditional GWAS for schizophrenia and bipolar disorder (affected vs. controls) but also performed GWAS for schizophrenia and bipolar disorder compared to controls and schizophrenia compared to bipolar disorder (Bipolar Disorder and Schizophrenia Working Group of the Psychiatric Genomics Consortium 2018). These unique comparisons allowed us to use S-LDSC to pinpoint what cell populations may be modulating disease differences.

We first looked at how the “Schizophrenia vs. controls” GWAS compared to our results from the CLOZUK study. While the “Schizophrenia vs. controls” samples were also included in the CLOZUK study, the studies were performed independently, and thus show slightly different results (Figure 4A, 4B; Table S6). 13/27 cell populations are found to be significant in both studies with all excitatory neuron populations reaching significance in both with layer V neurons being most enriched. Furthermore, both broadly defined inhibitory neuronal populations as well as inhibitory medium spiny neurons are found in both. However, the Psychiatric Genomics Consortium (PGC)-only study showed significant enrichment in embryonic forebrain dopaminergic neurons and inhibitory VIP neurons (Figure 4B) whereas summary statistics from the CLOZUK study did not. The PGC-only study also failed to detect enrichment in astrocytes and inhibitory PV neurons (Figure 4B).

Next, we looked at bipolar disorder and found similar results to schizophrenia. Namely, all excitatory neuron populations showed heritability enrichment with the highest enrichment being seen in the individual excitatory layers (Figure 4C; Table S6). Additionally, both broadly defined inhibitory neuron populations show enrichment, mirroring schizophrenia. In contrast, while subsets of cortical inhibitory neurons are found to be enriched in schizophrenia, none are found to be enriched in bipolar disorder (Figure 4C; Table S6). Perhaps most strikingly, the consistent, high enrichment of inhibitory medium spiny neurons in schizophrenia is absent in bipolar disorder, potentially pointing towards important biological differences (Figure 4C; Table S6). Furthermore, we analyze the combined schizophrenia and bipolar cohort and see significant enrichment in the same excitatory and inhibitory neurons. However, in the combined analysis, both embryonic dopaminergic populations also reach significance along with oligodendrocytes (Figure 4D; Table S6).

Finally, we analyze the schizophrenia versus bipolar disorder cohort. Only four cell populations reach trait-wide significance and all four are excitatory neurons including excitatory neurons from cortical layers II-III, IV, V, as well as the dentate gyrus (Figure 4E; Table S6). This result provides orthologous support to extensive work that has shown layer-specific neuronal differences between schizophrenia and bipolar disorder (Chana et al. 2003; Rajkowska et al. 2001; Benes et al. 2001) as well as differences in dentate gyrus neuronal maturation seen in these diseases (Yu et al. 2014). Overall, through the wealth of GWAS data available, we are able to begin to tease apart the complex relationship between schizophrenia and bipolar disorder.

### Statistical fine-mapping of 177 schizophrenia loci reveals complex biological hypotheses

Ultimately, our goal was to not only identify cell populations relevant to disease but to use that data to prioritize variants in schizophrenia loci. We incorporated significantly enriched open chromatin annotations into statistical fine-mapping of 177 independent schizophrenia loci. We extracted all common SNPs (minor allele frequency ≥ 0.01 in 1000 Genomes European data) in LD with 179 independent lead SNPs (*r*^2^ ≥ 0.1 in 1000 Genomes European data) from the CLOZUK schizophrenia GWAS (Pardiñas et al. 2018) (see Methods). Note that since we identified proxy SNPs from independent GWAS signals, some SNPs are fine-mapped independently in multiple loci. The summary statistics for all proxy SNPs were extracted from GWAS data and split into loci based on independent lead SNPs. Two loci were excluded (see Methods) resulting in a total of 177 independent schizophrenia loci.

We used the fine-mapping program, PAINTOR (Kichaev et al. 2014; Kichaev and Pasaniuc 2015; Kichaev et al. 2017) in order to incorporate annotation data. The open chromatin profiles for the 13 significant cell populations for the CLOZUK schizophrenia GWAS were merged into one annotation for use in fine-mapping (see Methods). Schizophrenia loci were fine-mapped with and without annotation using a Markov Chain Monte Carlo sampling algorithm without specifying the number of causal SNPs in each locus. In total, 62,994 unique SNPs were fine-mapped with an average of 370 SNPs per locus (Table S7). When combining results both with and without annotation, 1,512 SNPs in 166 loci reached a posterior inclusion probability (PIP) of ≥ 0.1, 82 SNPs in 56 loci reached a PIP ≥ 0.5, and 30 SNPs in 23 loci reached a PIP ≥ 0.9 (Table S7). All fine-mapping results can be found in Table S8.

We explored how all variants with a PIP ≥ 0.1 impact open chromatin regions in schizophrenia enriched cell populations. This cutoff was chosen because it has been shown to have a high benefit-to-cost ratio for follow-up experiments (Kichaev et al. 2014). We observed 281 unique SNPs across 104 loci achieve a PIP ≥ 0.1 and overlap with an open chromatin region present in at least one schizophrenia enriched cell population (Table S9). Furthermore, 242 SNPs are predicted to disrupt transcription factor binding sites as determined with ENCODE data (Table S10) with disruption of REST, EP300, and EGR1 TF motifs being the most common motifs disrupted (Table S11).

Since we used SNPs in LD with independent signals, not amalgamated loci, 2,317 SNPs were finemapped in more than one locus. 39 of those SNPs achieve a PIP ≥ 0.1 across multiple loci, potentially identifying the effects of single variants contributing to multiple reported “independent” signals (Table S12). One such example is rs11682175 which is in LD with two independent index SNPs located within the *VRK2* gene, rs75575209 (*r*^2^ = 0.105) and rs7596038 (*r*^2^ = 0.326) (Table S12). rs11682175 achieves high PIPs with annotation in both loci (0.999 and 0.864 in rs75575209 and rs7596038, respectively)(Table S12). While not reported in the CLOZUK GWAS, rs11682175 has been significantly associated with major depression (Wray et al. 2018), neuroticism (Nagel et al. 2018), and schizophrenia (Schizophrenia Working Group of the Psychiatric Genomics Consortium 2014). This SNP, however, does not intersect with any OCRs in the cell populations studied. We believe that we observe this result due to the “clumping” methodology used during the original GWAS (Pardiñas et al. 2018). While the top fine-mapped SNP (rs11682175) reaches genome-wide significance in the original GWAS, it seems to be grouped under rs7596038 during the first clumping procedure (based on LD and P-values) and was not reported as an “independent” signal (Pardiñas et al. 2018) (Figure S3). rs11682175 is much more significant than rs75575209, but since it is in low LD with rs75575209, it is included in that “independent” locus was well during our analysis. This high significance but low LD with the lead SNPs lead to this SNP (while not reported in the original GWAS) to rise to the top of both loci and may represent an independent signal masked by clumping.

For many loci, the hypotheses arising from our analyses are clear, consistent with known biology, and limited in scope; for others, the ongoing challenge is laid out in the breadth of SNPs highlighted by OCRs across a variety of cell types. We describe a few examples below.

The *GABBR2* locus is tagged by rs10985817 and contains 164 fine-mapped SNPs (Table S7). Two SNPs, rs10985817 and rs10985819, achieve a PIP ≥ 0.1. Both SNPs are encompassed by an OCR only present in dentate gyrus excitatory neurons (Table 1, Table S9) directly establishing a clear hypothesis. However, hypotheses emerging from many loci are not as immediately straightforward. One of the independent GWAS signals in an intron of *CACNA1C* (lead SNP rs2007044; 144 fine-mapped SNPs) contains eight SNPs that achieve a PIP ≥ 0.1 (Table 1; Table S7). Six of these SNPs intersect with at least one OCR in the populations studied with three intersecting exclusively with one excitatory neuron population (rs4765913, rs882195, rs11062170) (Table 1). rs4765913 resides exclusively in an OCR found in excitatory dentate gyrus neurons and has been significantly associated with bipolar disorder in multiple studies (Psychiatric GWAS Consortium Bipolar Disorder Working Group 2011; Charney et al. 2017). Further, one SNP, rs2239038, intersects OCRs found in excitatory neurons in multiple layers of the cortex (Table 1). Finally, two SNPs intersect with both excitatory and inhibitory neurons (rs1860002, rs2238057) (Table 1) indicating that *CACNA1C* expression may be modulated in both populations. Thus although biologically informed hypotheses may be directly developed in this way, they often remain multifaceted.

**Table 1.** Summary of prioritized SNPs in the rs10985817, rs2007044, and rs4144797 schizophrenia loci. All SNPs that achieved a PIP > 0.1 are included. Information about the prioritized SNPs in the table includes reference SNP ID (“RSID”), the lead independent SNP identified in Pardinas, et al. (Pardiñas et al. 2018) (“Lead SNP”), the PIP of each SNP when enriched annotations were not included (“PIP with no annotation”), the PIP of each SNP when enriched annotations were included (“PIP with annotation”), and the cell populations in which the variant intersects with open chromatin (“Cell populations”). Posterior inclusion probability, PIP; Single nucleotide polymorphism, SNP.

| RSID | Lead SNP | PIP with no annotation | PIP with annotation | Cell Populations |
| --- | --- | --- | --- | --- |
| rs10985817 | rs10985817 | 0.089 | 0.185 | Excitatory DG* |
| rs10985819 | rs10985817 | 0.067 | 0.146 | Excitatory DG* |
| rs4765913 | rs2007044 | 0.111 | 0.24 | Excitatory DG* |
| rs2007044 | rs2007044 | 0.342 | 0.206 | None |
| rs882195 | rs2007044 | 0.084 | 0.181 | Excitatory DG* |
| rs1860002 | rs2007044 | 0.089 | 0.18 | Inhibitory*, Excitatory Layers II-III, Excitatory Layer IV |
| rs2239038 | rs2007044 | 0.069 | 0.129 | Excitatory Layer V, Excitatory Layers II-III, Excitatory Layer IV |
| rs11062170 | rs2007044 | 0.055 | 0.117 | Excitatory Layer IV |
| rs2238057 | rs2007044 | 0.053 | 0.106 | Inhibitory VIP, Inhibitory Gad2, Excitatory Layers II-III |
| rs4298967 | rs2007044 | 0.117 | 0.094 | None |
| rs181813160 | rs4144797 | 0.906 | 0.974 | Neun neg, Embryonic DA Midbrain, Inhibitory VIP, Inhibitory*, Inhibitory MSN*, Inhibitory Gad2, Embryonic DA Forebrain, Excitatory Layer VI*, Excitatory Layer VI, Excitatory Layer V, Excitatory Layers II-V*, Excitatory Layers II-III, Excitatory Layer IV, Excitatory DG*, Excitatory Camk2a, Cones (green), Cones (blue), Astrocytes* |
| rs188020433 | rs4144797 | 0.942 | 0.967 | None |
| rs778371 | rs4144797 | 0.258 | 0.363 | Excitatory Layers II-V* |
| rs73102769 | rs4144797 | 0.159 | 0.204 | Inhibitory MSN*, Excitatory Layer V, Excitatory Layer IV |
| rs1878289 | rs4144797 | 0.139 | 0.195 | Embryonic DA Midbrain, Excitatory Layers II-V* |
| rs4144797 | rs4144797 | 0.082 | 0.189 | Rods, Neun neg, Embryonic DA Midbrain, Inhibitory VIP, Inhibitory Gad2, Embryonic DA Forebrain, Excitatory Layer VI, Excitatory Layer V, Excitatory Layers II-V*, Excitatory Layers II-III, Excitatory Layer IV, Excitatory DG*, Excitatory Camk2a, Cones (green), Cones (blue), CD8 T-cells, CD8 T-cells human, CD4 T-cells, CD4 T-cells human |
| rs2592127 | rs4144797 | 0.123 | 0.18 | Embryonic DA Midbrain, Excitatory Layer VI*, Excitatory Layer VI, Excitatory Layer V, Excitatory Layers II-V* |
| rs2675960 | rs4144797 | 0.076 | 0.115 | Embryonic DA Midbrain, Inhibitory VIP, Inhibitory MSN*, Inhibitory Gad2, Embryonic DA Forebrain, Excitatory Layer VI*, Excitatory Layer VI, Excitatory Layer V, Excitatory Layers II-V*, Excitatory Layers II-III, Excitatory Layer IV, Astrocytes* |
| rs4144795 | rs4144797 | 0.066 | 0.112 | Rods, Neun neg, Embryonic DA Midbrain, Microglia*, Inhibitory*, Inhibitory PV, Inhibitory Gad2, Embryonic DA Forebrain, Excitatory Layer VI, Excitatory Layer V, Excitatory Layers II-III, Excitatory Layer IV, CD8 T-cells, CD4 T-cells, CD4 T-cells human |
| rs1878287 | rs4144797 | 0.08 | 0.105 | Inhibitory*, Excitatory Layer VI*, Excitatory Layer VI, Excitatory Layer V, Excitatory Layers II-V* |
| rs938575 | rs4144797 | 0.112 | 0.068 | None |
| rs4973569 | rs4144797 | 0.103 | 0.053 | None |
| rs778363 | rs4144797 | 0.116 | 0.051 | None |

Like the *CACNA1C* locus, the locus tagged by rs4144797 contains 395 finemapped SNPs spread throughout a locus containing the *GIGYF2*, *KCNJ13*, *SNORC*, and *NGEF* genes on chromosome 2 (Figure 5A). This locus contains 13 SNPs that achieve a PIP ≥ 0.1 of which 9 intersect with an OCR from a variety of schizophrenia enriched cell populations (Table 1). Two of these SNPs are particularly interesting as they are located in promoter regions of genes. rs181813160 is located in the promoter of *NGEF* and the lead SNP, rs4144797, is located in the promoter of *GIGYF2* (Figure 5A). Both SNPs intersect with an OCR in a wide array of cell populations with rs181813160 intersecting 12/13 enriched populations and rs4144797 intersecting 8/13 enriched populations (Table 1). rs181813160 has the highest PIP (~0.97) in our dataset that also intersects an OCR, making it a prime causal candidate. Using ENCODE data, we find rs181813160 is predicted to strongly disrupt a potential binding site for 21 different transcription factors including immediate early genes, *EGR1*-*EGR4* (Figure 5B; Table S10). *EGR1* plays a major role in modulating synaptic plasticity and neuronal activity and EGR family members are down-regulated in schizophrenic brains (Yamada et al. 2007; Duclot and Kabbaj 2017). *NGEF* regulates the growth of axons and dendrites in neurons (Shamah et al. 2001; Blackmore et al. 2010), strengthening a hypothesis that would link this locus to synaptic dysfunction in schizophrenia (Fromer et al. 2014; Purcell et al. 2014; Sekar et al. 2016; Cannon 2015; Sellgren et al. 2019). Furthermore, rs4144797 is predicted to strongly create an EGR1 binding site (along with impacting 6 other TF binding sites), linking both promoter region variants (Figure 5B,Table S10).

**Figure 5.**
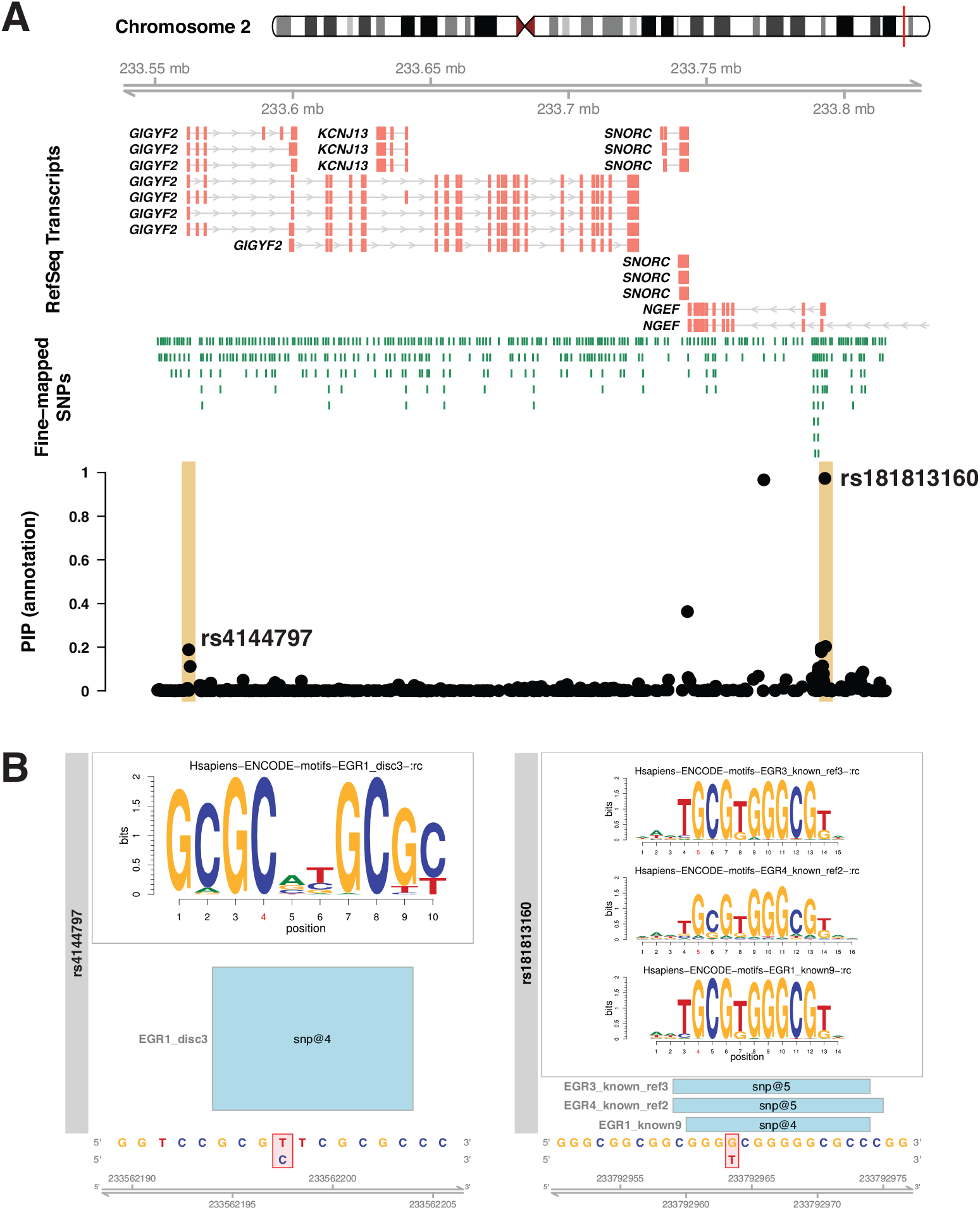
Fine-mapping prioritizes SNPs in the schizophrenia-associated locus surrounding the NGEF gene. A) A visualization of the schizophrenia-associated locus identified by the lead SNP, rs4144797. The plot displays the RefSeq transcripts and fine-mapped SNPs in the region in addition to the posterior inclusion probabilities (PIP) when annotation was included for all SNPs. The lead SNP (rs4144797) and the SNP with the highest PIP (rs181813160) are highlighted. B) EGR family motifs derived from ENCODE data that are created (rs4144797) or disrupted (rs18181360) in the rs4144797 locus. The nucleotides impacted are highlighted in red underneath the motif and named in the bars under the motif box.

## Discussion

Despite the capacity of GWAS to inform genetic architecture, connecting this to the risk, genesis and progression of disease has remained a stubborn challenge. This challenge is particularly stark in schizophrenia, where association of 179 independent loci implicates thousands of noncoding variants in disease risk without providing a systematic and biologically informed strategy to construct hypotheses. Genomic data is being increasingly used as a guide for the construction of hypotheses for common variant involvement in disease risk. Although a significant challenge, systematic functional testing of disease-associated, non-coding variation will be facilitated by the development of biologically informed hypotheses. This study emphasizes the value of strategies which seek cellular context for disease risk and non-coding variation as a prelude to massively parallel functional studies in cell types whose selection is truly biologically informed. It demonstrates the power of obtaining cellular surrogates from mice which may not be readily obtained from humans and opens the door to use the mouse model in the generation of temporal and sensitized cellular data to better inform human trait heritability and functional dissection.

Our study provides orthologous confirmation of the contribution of cortical and interneuron populations in neuropsychiatric disorders through chromatin data. In schizophrenia specifically, OCR signatures from cortical (both excitatory and inhibitory) populations are most enriched for schizophrenia heritability. We demonstrate a clear increase in the enrichment of heritability for schizophrenia from layer II-III, reaching a maximum at discrete layer V excitatory neurons and single-nuclei populations containing layer V neurons. Through the wealth of GWAS available, we were also able to illuminate cell specific differences and similarities between schizophrenia and bipolar disorder. Taking this data further, we prioritize schizophrenia variants through fine-mapping variants using open chromatin profiles from enriched cell populations. Inherently, these data establish a tableau of testable hypotheses that may be taken off the “shelf” into the lab environment. We identify SNPs in 104/177 tested schizophrenia-associated loci (~59%) that may now be considered candidates for functional testing in their specific cellular contexts. We see examples of relatively straightforward loci (*GABBR2* locus) and observations of more complicated hypotheses in which multiple variants within a risk haplotype may be exerting their effects, potentially simultaneously, in overlapping and unique cell populations (*CACNA1C* and *NGEF* loci). As a whole, these data take a critical next step in obtaining functional insight to common disease.

Although clearly powerful, the capacity to observe enrichment remains dependent on the availability of biological relevant datasets including cell types, developmental stages, or physiological states. The fact that these observations are facilitated by datasets generated in mice only serves to expand the potential application of this approach, reinforcing mice as a lens by which to study the genetics underlying common human phenotypes (Hook et al. 2018; McClymont et al. 2018). It establishes the feasibility of generating lineage-specific data where access in humans is more challenging. It makes available an almost limitless collection of cell populations with potential disease relevance; across the spectrum of developmental stages; in the presence or absence of genetic, chemical, and behavioral perturbation. Our data is consistent with recently published single-cell ATAC-seq work (Cusanovich et al. 2018). Here we demonstrate, although useful, single-cell acquisition is not necessary to achieve the cell layer-based resolution. Furthermore, the sparse nature of chromatin data obtained from individual cells necessitates sequencing large numbers of cells (Cusanovich et al. 2018) or additional information from RNA to optimize cell identification (Cao et al. 2018). Even with multiple levels of information, some cell populations delineated through single-cell assays cannot be reliably identified (Cusanovich et al. 2018; Cao et al. 2018; Preissl et al. 2018). Bulk assays, at this time, may thus prove more immediately feasible and flexible.

While we generate results that support previous studies, there are limitations to the breadth and depth of data we have assembled. We cannot arrive at conclusions about the disease relevance of any cell-type that was not tested here. Additionally, open chromatin profiles lack the biological interpretation provided by histone marks when trying to identify functional regulatory DNA. We expect these issues to be solved progressively by increasing the resolution, quality, and completeness of chromatin and histone data in tandem with decreasing cell numbers needed to obtain high quality data.

Further, our results rely on lifting over mouse data to the human genome. While we optimize lifting over and show the vast majority of mouse peaks have a human syntenic ortholog, the entire landscape of regulatory DNA present in these cell populations in humans cannot be queried. Although the extent to which this limits immediate progress is unclear, we show on average, 78% of mouse-derived human peaks are open in humans and that these profiles recapitulate heritability enrichment results from a variety of phenotypes. This gives us confidence in our approach, however, efforts to obtain single-cell data from human brain samples make future human datasets a possibility (Lake et al. 2018).

Finally, our analysis focuses on common SNPs due to the underlying model used in S-LDSC (Finucane et al. 2018). We, therefore, cannot come to any conclusions about the contribution of rare variation. This may cause the exclusion of true causal variants. We acknowledge this may be compounded by a focus on SNPs. Other more complex variation has been shown to be important in schizophrenia loci and cannot yet be fully assayed with this method (Sekar et al. 2016; Song et al. 2018).

Overall, our data define a spectrum of immediately testable hypotheses, implicating specific variants as potentially modulating the activity of cis-regulatory elements in discrete cellular contexts across phenotypes. Taken collectively the capacity to move directly from GWAS to design of functional tests by using mouse-derived data represents a significant step forward in the dissection of common human phenotypes.

## Methods

### Obtaining ATAC-seq data

Raw ATAC-seq sequencing data was primarily obtained from the Gene Expression Omnibus (GEO) except single-nuclei ATAC-seq data (Preissl et al. 2018), which was obtained from the author’s website (URLs). All details about the downloaded sequencing data can be found in Table S1.

Additional steps were needed in order to aggregate ATAC-seq reads from individual nuclei into cell populations. A list of barcode names (cells) and the clusters to which they belonged in Preissl, et. al., were obtained from the authors of the original paper (personal communication). Crucially, these barcodes were included in the name of each sequencing read. Barcodes were grouped according to their cluster identity and paired-end reads belonging to each cell population were extracted via sequencing read name using the BBMap script ‘demuxbyname.sh’ with the parameters ‘substringmode’ (URLs). FASTQ files for each barcode in each replicate were then combined into a single FASTQ file for each cluster. This method had the advantage of only extracting reads originating from cells that had passed quality control measures (Preissl et al. 2018).

### Alignment and peak calling

Paired-end reads were aligned to the mouse genome (mm10/GRCm38; URLs) using bowtie2 (version 2.2.5; URLs) (Langmead and Salzberg 2012) with the following parameters: ‘-p 15 --local -X2000’. Paired-end reads aligning to the mitochondrial genome as well as random and unknown chromosomes were removed. SAMtools (Li et al. 2009) was used to remove duplicate reads (v0.1.9), improperly paired reads (v1.3.1), and reads with a mapping quality score of less than or equal to 30 (v1.3.1).

Replicates for each cell population were then merged into a single bam file and peak summits were called for each mouse cell population (25 in total) using the MACS2 (v. 2.1.1.20160309) (Zhang et al. 2008) ‘macs2 callpeak’ function with the following parameters: ‘--seed 24 --nomodel --nolambda --call-summits --shift −100 --extsize 200 --keepdup all --gsize mm’. Regions that are considered artifacts of ATAC-seq and other chromatin assays in mm10 (so called ‘blacklist regions’; see URLs) were removed using BEDtools (v2.27.0) ‘intersect’ (Quinlan and Hall 2010). Raw MACS2 output can be found on Zenodo (see Data Access).

In order to perform downstream comparisons with data generated in mice, raw ATACseq data from CD4 and CD8 T-cells were obtained (Corces et al. 2016) (Table S1). These data were processed in the same manner as the mouse data above with the exception that the reads were aligned to the human genome (hg19).

Since we were comparing ATAC-seq datasets with vastly different sequencing depths and numbers of called summits, we applied a recently introduced filtering strategy for ATAC-seq peaks (Corces et al. 2018). For each dataset, we summed the MACS2 peak scores and divided that number by one million (total score per million). We then divided each individual peak score by the total score per million for that dataset to produce a “score per million” (Corces et al. 2018). Ultimately we chose a “score per million” cutoff of two as that would equate to a P-value per million of 0.01.

### Relationship between public mouse sets

Summits called in each population were made into uniform 501 bp peaks by adding 250 bp to each side of the summit. Peaks were then merged into a union set of peaks using BEDtools ‘merge’ with default parameters. This final set of filtered and merged peaks contained a total of 433,555 peaks.

In order to obtain a count matrix for cell population comparison, featureCounts (v1.6.1) was used (Liao et al. 2014). First, the union set of ATAC-seq peaks was manually converted to an SAF file (see Subread website; URLs). The command ‘featureCounts’ was used with the ‘-T 10 -F SAF’ parameters in order to obtain a count matrix. BEDtools ‘nuc’ was used with a FASTA file of combined mm10 chromosome sequences obtained from the UCSC Genome browser (URLs) in order to calculate GC content for each peak.

The count matrix, the count matrix summary file, and the peak GC content file were read into the R statistical environment (URLs). Data were transformed into log2(count + 1) counts and the CQN R package (Hansen et al. 2012) and ComBat from the SVA R package (Leek et al. 2012) were used to quantile normalize counts and correct CQN normalized counts for type of experiment (single-nuclei or bulk). Principal component analysis (PCA) was performed using all peak counts with the R functions ‘prcomp()’ with default settings and “scale. = TRUE” setting. t-SNE was performed with the first 6 principal components from PCA using the ‘tsne’ package in R with the ‘tsne()’ function with the following parameters: perplexity = 5, max_iter = 10000, whiten = T. Additionally, the Pearson correlation between corrected peak counts was used to hierarchical cluster the data. Correlations were converted to distances by subtracting the absolute value of the correlations from 1. Clustering was performed using the R function ‘hclust’ with ‘method = “ward.D2”’ and figures were produced with custom R scripts.

### Liftover

All strategies used the lift over script “bnMapper. py” from the bx-python software package (Denas et al. 2015) (URLs) along with the “reciprocal best” mm10 to hg19 chain file (mm10.hg19.rbest.chain. gz) from UCSC genome browser (URLs). Three different lift over strategies were compared: one using the called summits and two using the uniform, unmerged 501 bp peaks. The first strategy lifted over the single bp summit sets with the settings: ‘-f BED12’. The second strategy lifted over the 501 bp peak sets again with the settings: ‘-f BED12’. The third strategy again used the 501 bp peaks with the settings: ‘-f BED12 -g 20 -t 0.1’. This third strategy has been employed previously (Vierstra et al. 2014) and it applies a more strict lift over which limits the size of the gaps allowed in the mapped sequences. Ultimately, the lift over of the peak summits was used for all subsequent analyses. After lift over to hg19, 250 bp was added on to both sides of each summit to create peaks. Overlapping peaks for each annotation were merged using BEDtools ‘merge’ with default parameters. Human regions that are blacklisted either by the ENCODE consortium or ATAC-seq users were removed (URLs).

### Comparison to publicly available human open chromatin data

Human open chromatin profiles derived from mouse data were compared to imputed Roadmap Epigenetic Project DNase I hypersensitivity data from 127 human tissues and cell populations (Ernst and Kellis 2015) and ATAC-seq data from neurons isolated from 14 human brain regions (Fullard et al. 2018) (URLs). Comparisons were made using the BEDtools ‘jaccard’ command with default parameters. Overlaps were calculated for each annotation (Table S4).

### Partitioning heritability with linkage disequilibrium score regression (S-LDSC)

All necessary components needed to run S-LDSC including baseline scores, PLINK files, frequency files, weights, and SNPs, were downloaded from the Broad Institute (URLs; Table S13). All files were ‘1000G_Phase3’ versions. Additionally, Roadmap Epigenetic Project LDSC files were used as additions to the baseline model as was done in a previous application of LDSC on ATACseq data (Finucane et al. 2018). These were also obtained from the Broad Institute (URLs; Table S13).

Summary statistics for 64 GWAS were obtained from a variety of sources as either “raw” summary statistics or summary statistics that were pre-processed in the LDSC pipeline (Table S5; URLs). 48 of the summary statistics were obtained from the Alkes Price group as either preprocessed, published summary statistics or “raw” summary statistics of UK Biobank phenotypes (Table S5; URLs). 16 of the summary statistics were handpicked and were mostly from neurological traits including schizophrenia. “Raw” GWAS summary statistics were downloaded and processed using the ‘munge_sumstats.py’ script (LDSC v1.0.0). Specific command parameters used to process the data are listed in Table S5. Note, processed summary statistics from the CLOZUK schizophrenia GWAS (Pardiñas et al. 2018) needed minor modifications after processing (Table S5).

Annotation files needed for analysis were created using the ‘make_annot.py’ script included in the LDSC software (v1.0.0; URLs) while specifying the following parameters: --bed-file; --bimfile; --annot-file. LD score files needed for analysis were created with the ‘ldsc.py’ script with the following parameters: --l2; --bfile; --ld-wind-cm 1; --thin-annot; --annot; --out; --print-snps. Cell-type partitioned heritability calculations (also referred to as S-LDSC) were performed with the ‘ldsc.py’ script with the following parameters: --h2-cts; --ref-ld-chr; --ref-ld-chr-cts; --w-ld-chr.

The P-values for heritability enrichment are based on a one-sided test for the regression coefficient being greater than 0. This allowed for a direct comparison of the magnitude of enrichment (i.e. higher P-value = higher enrichment). For more information, see Finucane, et al., 2015 (Finucane et al. 2015) and LDSC website (URLs). Partitioned heritability calculations for all traits were combined and analyzed in R. The creation of plots was carried out using custom R scripts. The level of significance was set for LDSC results as the Bonferroni corrected P-value when taking into account all summary statistics and cell populations tested (0.05/(27*64) = 0.00002894; -log10(P) = 4.53857).

### Fine-mapping SNPs in schizophrenia loci

#### Finding proxy SNPs

A total of 179 genome-wide significant, independent index SNPs from the CLOZUK SZ study were obtained (Pardiñas et al. 2018). In order to assay all common SNPs within LD of the index SNPs, the function ‘get_proxies’ from the R package ‘proxysnps’ (URLs) was used with the following parameters: window_size = 2e6, pop = “EUR”. Only SNPs with an *r*^2^ ≥ 0.1 from an index SNP and a minor allele frequency (MAF) ≥ 1% were retained for fine-mapping.

This method obtained 71,344 unique SNPs with reference SNP (RS) numbers across 177 loci. Note that some SNPs are shared between loci. The index SNP as reported in Pardinas, et al., could not be used to identify proxy snps in seven loci for various reasons. Instead a suitable replacement was used based on LD or on changes in SNP databases over time (Table S14). In addition, two genome-wide significant loci were excluded from the analysis. The locus with the index SNP rs1023497 was excluded because it is not a biallelic variant in 1000 Genomes data so proxies were not found. The second locus was the MHC locus (rs3130820) because of its complicated LD structure and because it is generally excluded from LDSC analysis (Finucane et al. 2018).

#### File setup for fine-mapping

In order to discover disease-relevant variants within each locus, we statistically fine-mapped all 177 SZ loci using PAINTOR (v3.1; URLs)(Kichaev et al. 2014; Kichaev and Pasaniuc 2015; Kichaev et al. 2017). PAINTOR was chosen for its ability to use summary statistics, run simulations on multiple loci at once, and incorporate chromatin annotation data. Proxy SNPs were merged with summary statistics from Pardinas, et al. (Pardiñas et al. 2018) (URLs) leaving 62,994 unique SNPs. The merged data was then split into 177 loci based on the index SNP and formatted for use in PAINTOR by using custom R scripts. The number of SNPs in each locus ranged from 7 to 1919 (Table S15). The loci were used to create both the LD and annotation files needed to run PAINTOR.

LD files were created with the script ‘CalcLD_1KG_VCF.py’ included in PAINTOR with the following parameters: --reference; --mapl; --effect_allele A1; --alt_allele A2; --population EUR; --Zhead Zscore; --position pos. The 1000 Genomes reference VCF used was imputed and filtered by Beagle (Browning et al. 2018) since the program used to find proxy SNPs (‘proxysnps’) used the same VCF (URLs). Note that the downloaded ‘CalcLD_1KG_VCF.py’ script was modified as suggested on the PAINTOR GitHub page so if the Z-score was flipped when calculating LD, the alleles were also flipped (URLs). It was also modified so ambiguous SNPs would not be removed.

Annotation files were created using the ‘AnnotateLocus.py’ script included with PAINTOR with the following key parameters: --chr chr --pos pos. Python syntax in this script was modified in order for it to run (see Github). As suggested by the PAINTOR authors, the correlations between annotations found to be significant in LDSC were calculated using custom R scripts. All significant annotations had a Pearson correlation > 0.2 (the cut-off suggested by authors), so all annotations were merged. Annotation files for all loci were reproduced with the merged annotation.

#### Running PAINTOR fine-mapping

In order to reduce the time and computational burden required to estimate annotation enrichments in PAINTOR and perform the fine-mapping with sufficiently long Monte Carlo Markov Chain (MCMC) simulation, the merged annotation enrichment was estimated with a shorter MCMC with the following key parameters: -mcmc; -burn_in 5000; -max_samples 1000 -num_chains 5; -set_seed 3; -MI 30. The enrichment estimates for the baseline model and the annotation model were then used in subsequent analyses.

In order to perform robust fine-mapping using MCMC, enrichments estimated above were used as input to a fine-mapping strategy using PAINTOR MCMC simulations with the following key parameters: -mcmc; -burn_in 100000; -max_samples 1000000 -num_chains 5; -set_seed 3; -MI 1. Fine-mapping was run both with and without merged annotations with the parameter for supplying enrichment estimates set at ‘-gamma_intial 3.79521’ for the no annotation simulation and ‘-gamma_initial 3.79521, −0.939523’ set for the simulation including annotation. The number of samples used for ‘-burn_in’ and ‘-max_samples’ parameters were chosen based on parameters set for MCMC fine-mapping with other methods (Banerjee et al. 2018). Visualizations of fine-mapping results were created with custom R scripts.

#### SNP transcription factor binding site disruption

In order to explore the functional impact of finemapped SNPs on transcription factor binding sites, the R program motifbreakR was used (Coetzee et al. 2015). All SNPs with a PIP ≥ 0.1 in either of the fine-mapping simulations (with or without annotations) and overlap with an open chromatin region from a S-LDSC schizophrenia enriched cell-population were used. The following parameters were used in the ‘snps.from.rsid()’ function from motifbreakR in order to analyze the variants: ‘dbSNP = SNPlocs. Hsapiens.dbSNP144.GRCh37; search.genome = BSgenome.Hsapiens.UCSC.hg19’. SNPs were then scanned for modification of transcription factor binding sites as defined by ENCODE by using the ‘motifbreakR()’ function with the following parameters: filterp = TRUE; pwmList = encodemotif; threshold = 1e-4; method = “ic”; bkg = c(A=0.25, C=0.25, G=0.25, T=0.25); BPPARAM = BiocParallel::bpparam().

### URLs

#### Preissl, et al. full data set

http://renlab.sdsc.edu/r3fang/snATAC/

#### mm10 Bowtie2 index

ftp://ftp.ccb.jhu.edu/pub/data/bowtie2_indexes/mm10.zip

#### Bowtie2

http://bowtie-bio.sourceforge.net/bowtie2/index.shtml

#### Subread FeatureCounts

http://bioinf.wehi.edu.au/featureCounts/

#### Broad Institute LDSC summary statistics files

https://data.broadinstitute.org/alkesgroup/sum-stats_formatted/; https://data.broadinstitute.org/alkesgroup/UKBB/

#### LDSC

https://github.com/bulik/ldsc

#### BBMap

https://sourceforge.net/projects/bbmap/

#### R statistical software

http://www.r-project.org/

#### tsne R package

https://github.com/jdonaldson/rtsne

#### mm10 fasta sequence

http://hgdownload.cse.ucsc.edu/goldenPath/mm10/chromosomes/

#### ENCODE blacklisted regions

http://mitra.stanford.edu/kundaje/akundaje/release/blacklists/ (Downloaded May 4, 2018)

mm10: mm10.blacklist.bed.gz

hg19: wgEncodeHg19ConsensusSignalArti factRegions.bed.gz

#### ATAC-seq mitochondrial blacklisted regions

https://sites.google.com/site/atacseqpublic/atac-seq-anal-ysis-methods/mitochondrialblacklists-1 (Downloaded: May 4, 2018)

mm10: JDB_BLACKLIST.MM10.BED hg19: JDB_BLACKLIST.HG19..BED

#### Bx-python

https://github.com/bxlab/bx-python

#### Reciprocal best liftover chain

https://hgdown-load-test.gi.ucsc.edu/goldenPath/hg19/vsMm10/re-ciprocalBest/

#### Roadmap DNase I imputed data

https://egg2.wustl.edu/roadmap/data/byFileType/peaks/consolidated-Imputed/narrowPeak/

#### BOCA ATAC-seq

https://bendlj01.u.hpc.mssm.edu/multireg/resources/boca_peaks.zip

#### proxysnps

https://github.com/slowkow/proxysnps

#### PAINTOR

https://github.com/gkichaev/PAINTOR_V3.0

#### PAINTOR modification

https://github.com/gkichaev/PAINTOR_V3.0/issues/17

#### BEAGLE VCF

http://bochet.gcc.biostat.washing-ton.edu/beagle/1000_Genomes_phase3_v5a/

### Data Access

The sources for publicly available ATAC-seq data can be found in Table S1 and are described in the Methods. Documentation of code is available on GitHub (https://github.com/pwh124/open_chromatin). Access to data including peaks and all files for heritability enrichment analyses and fine-mapping are available viaZenodo(https://doi.org/10.5281/zenodo.3253181).

## Acknowledgements

We would like to give special thanks to Sebastian Preissl, David U. Gorkin, and Rongxin Fang, for providing the information necessary to process single-nuclei ATAC-seq data. Funding: This research undertaken at Johns Hopkins University School of Medicine was supported in part by awards from NIH (NS62972 and MH106522) to ASM. Author contributions: PWH and ASM designed the study and wrote the manuscript. PWH implemented the computational algorithms to process the raw data and conduct analyses. PWH and ASM analyzed and interpreted the resulting data. PWH contributed novel computational pipeline development. Correspondence to ASM. Competing interests: The authors declare no competing interests.

---

**Figure S1.**
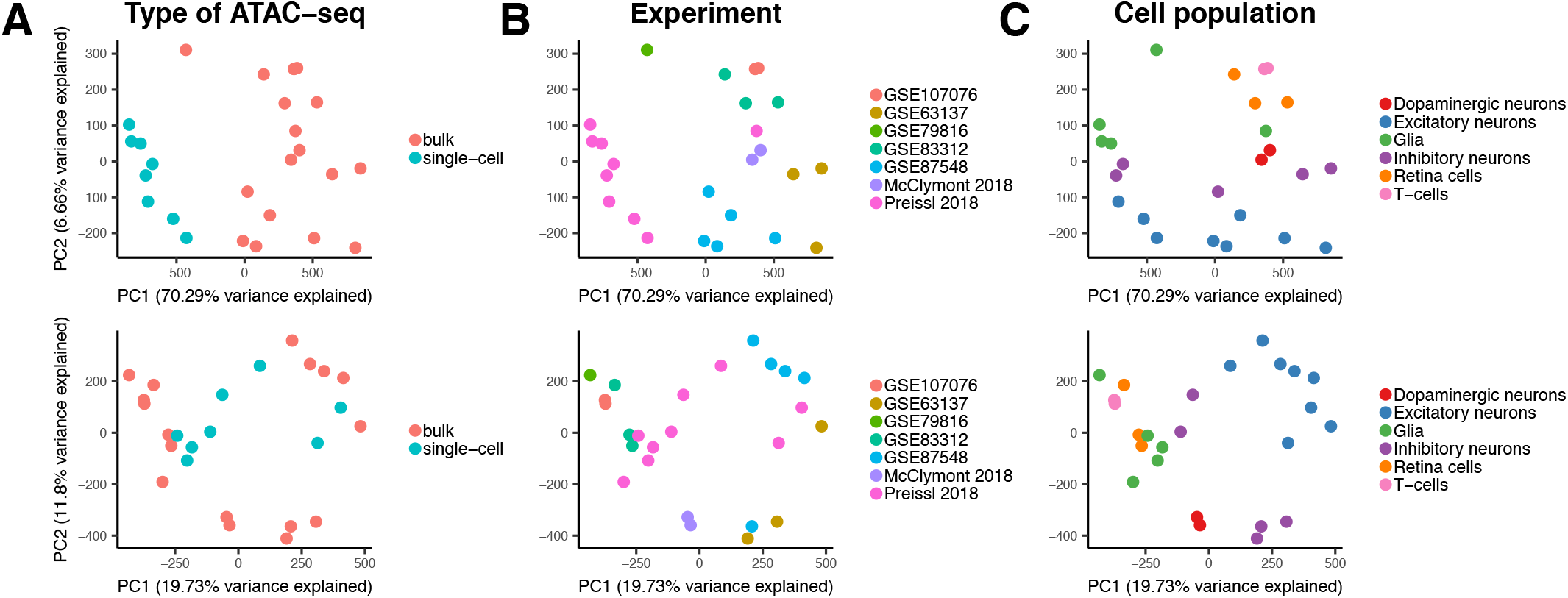
Summary of principal component analysis of ATAC-seq cell population peak read counts. A, B, C) PC1 vs. PC2 for log2(counts + 1) (top) and quantile normalized and batch corrected log2(counts + 1) (bottom). Cell populations are colored according to type of ATAC-seq performed (“bulk” or “single-cell”), B) experiment from which they came, and C) broad cell population category. All meta-information can be found in Table S1.

**Figure S2.**
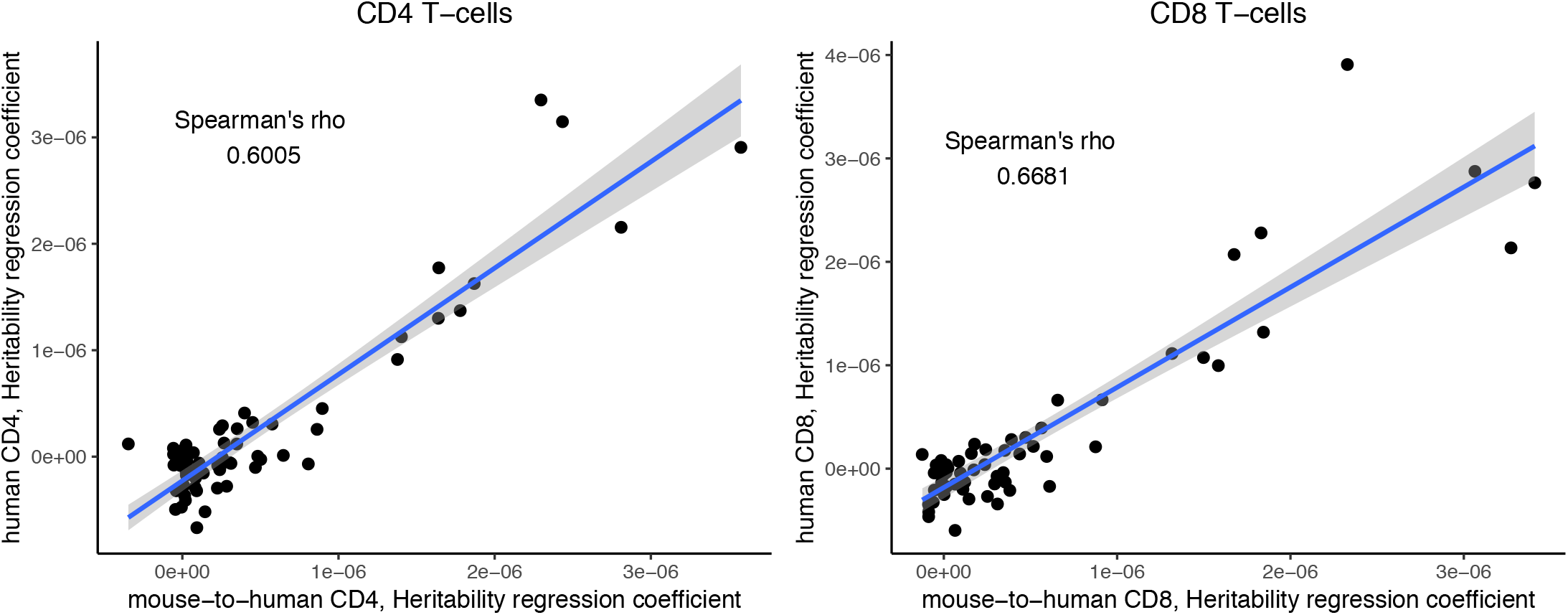
S-LDSC results are correlated between human T-cell open chromatin profiles and mouse-derived human open chromatin profiles. A, B) Scatterplots of S-LDSC heritability regression coefficients between orthologous cell populations of A) CD4 and B) CD8 T-cells. Both plots show a linear model fit to the data with a 95% confidence interval.

**Figure S3.**
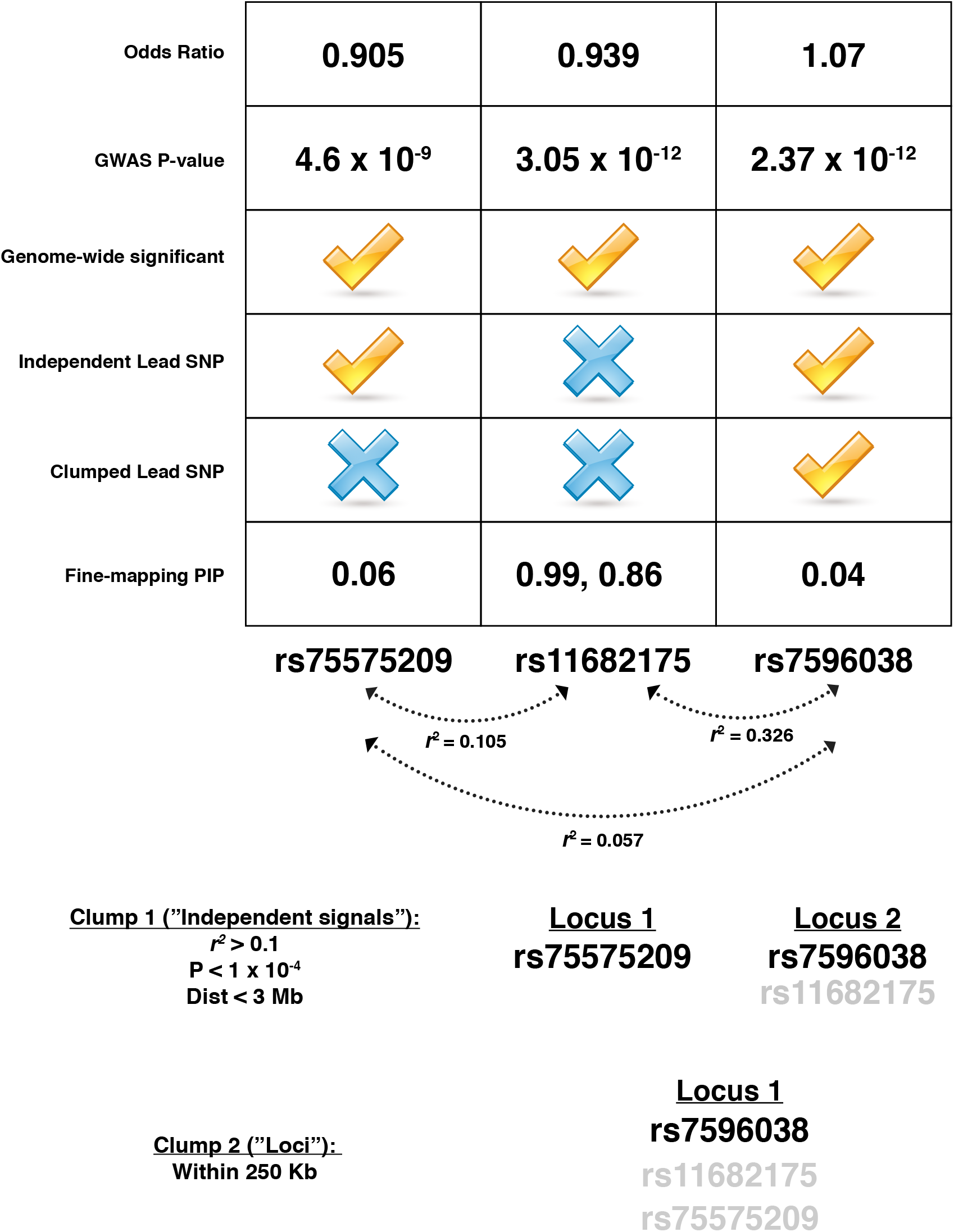
rs11682175 is the top fine-mapped SNP in the rs75575209 and rs7596038 independent schizophrenia associated loci. Summary statistics (“Odds Ratio” and “GWAS P-value” are shown at the top of the figure for all three SNPS (rs11682175 and lead SNPs). The boxes below the summary statistics show a checkmark if the SNPs meet that criteria in Pardiñas et al. 2018. These criteria include whether the SNP was genome-wide significant in the original study, whether it was reported as an independent lead SNP, whether it was reported as a clumped lead SNP and what the fine-mapping posterior probabilities including annotations were for each SNP. Below the chart are the LD relationships between SNPs as calculated in 1000 Genomes European data (Phase 3). The bottom of the figure shows how the clumping procedures used in the original study lead to three genome-wide SNPs being combined into two independent lead SNPs and one reported locus.

## Supplemental Material

***Supplemental Table S1. Description of all publicly available ATAC-seq used in this study***. Included is the cell population name used in the paper, the file name used for the cell population data, a description of the cell population, the type of ATAC-seq, a broader cell type classification of cell populations, the Pubmed or bioRxiv ID for the publication of the data, source of the data, and any associated file names.

***Supplemental Table S2. Summary of mouse peak data***. This includes the cell population name, the number of summits called in the mouse genome (“mm10_ summits”), the number of summits that passed filtering (“mm10_filtered_summits”), and the number of peaks that resulted from the called summits (“mm10_peaks”).

***Supplemental Table S3. Summary of T-cell peak overlap data***. This includes the mouse-derived human open chromatin profile file name (“human. mouse filename”), the total number of mouse-derived human peaks (“total.peaks”), the filename of the human open chromatin data (“human filename”), the sample name used for the human data (“sample.name”) and the number of peaks that overlap between the two files (“overlap.count”).

***Supplemental Table S4. Summary of overlap data for all cell populations***. This includes the cell population name (“population”), the number and percentage of peaks that overlap with all Roadmap Epigenome Atlas data (“roadmap_peaks” and “roadmap_percent”), the number and percentage of peaks that overlap with brain related tissues in Roadmap Epigenome Atlas Data (“roadmap_brain_ peaks” and “roadmap_brain_percent”), the number and percentage of peaks that overlap with ATAC-seq from BOCA(Fullard et al. 2018) (“boca_peaks” and “boca_percent”), and the number and percentage of peaks that overlap with all the data combined (“combined_peaks” and “combined_percent”).

***Supplemental Table S5. Summary of all the summary statistics analyzed with S-LDSC***. This includes the phenotype name (“Phenotype”), the name of the source of the data (“Source”), the URL from which the data was downloaded (“URL”), PubMed ID for any accompanying publications (“PMID”), the filename of the summary statistics (“Filename”), the exact LDSC munge command used to process the data (“Munge_command”), and any additional notes.

***Supplemental Table S6. The results from S-LDSC for all 64 traits analyzed***. Each tab in the document is a different set of summary statistics. Each tab contains the cell population (“cell.population”), the broad type of cell (“type”), the phenotype analyzed (“pheno”), the heritability regression coefficient (“Coefficient”), the standard error of the heritability regression coefficient (“Coefficient_std_error”), the P-value of the coefficient (“Coefficient_P_value”), the -log10(P-value) (“p.log10”), and whether or not the cell population achieves significance (“signif.all”).

***Supplemental Table S7. Summary of 177 fine-mapped schizophrenia loci***. This contains the lead SNP identified in Pardinas, *et al*.(Pardiñas et al. 2018) (“lead.snp”), the total number of SNPs fine-mapped for each locus (“total.snps”), the number of SNPs that reach a PIP ≥ 0.1 (“≥ 0.1”), the number of SNPs that reach a PIP ≥ 0.5 (“≥ 0.5”), the number of SNPs that reach a PIP ≥ 0.9 (“≥ 0.9”), the number of SNPs that both reach a 0.1 PIP and overlap with an open chromatin region in an enriched cell population (“overlap.ten.enrich”), the number of SNPs that both reach a 0.5 PIP and overlap with an open chromatin region in an enriched cell population (“overlap.fifty. enrich”), the number of SNPs that both reach a 0.9 PIP and overlap with an open chromatin region in an enriched cell population (“overlap.ninety. enrich”), and the total number of causal variants calculated per locus by adding PIPs for both fine-mapping without annotation (“total.pp.null”) and with enriched annotation (“total.pp.annotation”).

***Supplemental Table S8. All results from finemapping 177 schizophrenia loci***. This includes the SNP ID (“id”), the chromosome (“chr”), the position (“pos”), the reference SNP ID (“rsid”), the A1 allele (“A1”), the A2 allele (“A2), the Z-score for schizophrenia (“Zscore”), the lead SNP (“lead.snp”), the R^2^ between the proxy SNP and the lead SNP (“r. squared”), the -log10(P-value) for the SNP in the schizophrenia GWAS (“-log10(P)”), the PIP when annotation was not included (“PIP_null”), and the PIP when the annotation was included (“PIP_anno”).

***Supplemental Table S9. All results from SNPs that achieve a PIP ≥ 0.1 and overlap with an open chromatin region from an enriched cell population***. All information mentioned in Table S8 are present. In addition, included in “binary matrix” indicating whether or not the SNP intersects open chromatin in that cell population (0 = “no”, 1 = “yes”). Note that instead of asterisks, single-nuclei datasets are indicated with the “_sc” suffix. Finally, the total number of cell populations that the SNP intersects with (“all.sum”) and the total number of enriched cell populations that the SNP intersects with (“enrich.sum”) are included.

***Supplemental Table S10. All results from motifbreakR***. This includes the name of the SNP (“rsid”), the lead SNP or SNPs it is associated with (“lead.snps”), the effect of the motif disruption (“effect”), the gene symbol of the transcription factor whose motif is disrupted (“geneSymbol”), the source of the motif data (“dataSource”), the name of the motif (“providerName” and “providerID”), and the sequence that is matched for the motif (“seqMatch”).

***Supplemental Table S11. A summary of how frequently the motif of a transcription factor is impacted by SNPs with a PIP ≥ 0.1***. Included is the name of the transcription factor (“TF.gene”) and the number of times it is disrupted by a SNP (“Freq”).

***Supplemental Table S12. All results of SNPs that achieve a PIP ≥ 0.1 in multiple schizophrenia loci***. All columns included in Table S8 are present.

***Supplemental Table S13. A summary of LDSC file downloads***. Includes file’s purpose (“LDSC files downloaded”) and the download link (“URL”).

***Supplemental Table S14. A summary of the SNPs used to extract proxy SNPs for schizophrenia loci***. This includes chromosome (“chr1”), start and of locus (“start” and “end”), the lead SNP for the locus (“lead. snp”), the SNP used to extract proxy SNPs (“search. snp”), and any notes about the search SNP (“notes”).

***Supplemental Table S15. A summary of the number of SNPs in each schizophrenia locus throughout the process of creating files to be fine-mapped by PAINTOR***. This includes the lead SNP (“index.snp”), the total number of proxies extracted (“all.proxies”) and the total number of proxies after merging with summary statistics (“snps.after.merge”).

